# High-resolution genetic and phenotypic analysis of *KIR2DL1* alleles and their association with pre-eclampsia

**DOI:** 10.1101/330803

**Authors:** Oisin Huhn, Olympe Chazara, Martin Ivarsson, Christelle Retiere, Tim Venkatesan, Hormas Ghadially, Ashley Moffett, Andrew Sharkey, Francesco Colucci

## Abstract

Killer-cell immunoglobulin-like receptor (KIR) genes are inherited as haplotypes, expressed by NK cells and linked to outcomes of infectious diseases and pregnancy. Understanding how genotype relates to phenotype is difficult because of the extensive diversity of the KIR family. Indeed, high-resolution KIR genotyping and phenotyping in single NK cells in the context of disease association is lacking. Here, we describe a new method to separate NK cells expressing allotypes of the KIR2DL1 gene carried by the KIR A haplotype (KIR2DL1A) from those expressing KIR2DL1 alleles carried by the KIR B haplotype (KIR2DL1B). We find that in KIR AB heterozygous individuals, different KIR2DL1 allotypes can be detected both in peripheral blood and in uterine single NK cells. Using this new method, we demonstrate that both blood and uterine NK cells co-dominantly express KIR2DL1A and KIR2DL1B allotypes, but with a predominance of KIR2DL1A variants, which associate with enhanced NK cell function. In a case-control study of pre-eclampsia, we show that *KIR2DL1A*, not *KIR2DL1B*, associates with increased disease risk. This method will facilitate our understanding of how individual *KIR2DL1* allelic variants affect NK cell function and contribute to disease risk.

## INTRODUCTION

Natural Killer (NK) cells are important effectors in immune responses to viruses and tumours (Kärre, Ljunggren, Piontek, & Kiessling, 1986). Increasingly, tissue resident NK cells are described which defy the classical paradigm of NK cells as ‘killers’ (Reviewed in (Björkström, Ljunggren, & Michaëlsson, 2016)). Rather, they fulfil tissue specific roles such as interacting with placental cells during early pregnancy (Hanna et al., 2006). NK cell function is tightly controlled by continuously integrating signals from activating and inhibitory receptors including the Killer-cell immunoglobulin-like receptors (KIR). This polymorphic gene family is expressed predominantly by NK cells and to a lesser extent T cells. Both activating and inhibitory KIR exist, some of which bind HLA class I ligands to regulate NK functional responses. In addition, inhibitory KIR that bind cognate HLA class I ligands act with other inhibitory receptors such as CD94/NKG2A, to tune the reactive potential of NK cells to their environment; a dynamic process known as education (Goodridge, Önfelt, & Malmberg, 2015). Because both KIR and HLA are highly polymorphic, this diversity creates a spectrum of NK cell function both within and between individuals. More than 200 studies demonstrate disease association to variants of KIR and their HLA ligands (Reviewed in (Béziat, Hilton, Norman, & Traherne, 2016)). These include associations with outcomes in infectious diseases such as HIV/AIDS (Martin et al., 2002; 2007) and hepatitis (Khakoo et al., 2004), complications of pregnancy such as pre-eclampsia (Hiby et al., 2010; 2004; Nakimuli et al., 2015) and the outcome of immunotherapy in cancer patients (Forlenza et al., 2016). However, the complexity of both the *KIR* and *HLA* gene families presents an obstacle to understanding how these genetic associations contribute to pathogenesis.

The diversity of the KIR system arises from several sources. Firstly, *KIR* genes are inherited as 2 haplotypes, classified based on gene content. The *KIR A* haplotype has fewer genes with mainly inhibitory KIR, whereas *KIR B* haplotypes contain additional activating *KIR*. In this way, individuals will differ by the restricted suite of *KIR* genes they possess. This is compounded by the fact that the same *KIR* gene can be found on multiple haplotypes. Therefore, copy numbers of specific *KIR* genes can also vary. A further source of variation is the extensive *KIR* allelic polymorphism, for example over 80 alleles have been described for *KIR3DL1/S1* (IPD-KIR database). In addition to the genetic diversity, KIR expression is stochastically regulated by two opposing sense and antisense promoters (Anderson, 2014). Thus, in a population of NK cells there is variegated expression of KIR or none at all. Moreover, *KIR* genes are located within the leukocyte receptor complex on chromosome 19 and segregate independently from their HLA ligands encoded on chromosome 6. As a result, individuals may not have all the cognate ligands for their particular KIR repertoire; this will affect the education status of their NK cells. The polymorphic nature of the KIR/HLA system complicates linking genotype to phenotype in the context of disease association studies.

The inhibitory receptor KIR2DL1 recognises HLA-C allotypes bearing a C2 epitope (C2^+^HLA-C), defined by lysine at position 80. A number of disease associations studies have linked KIR2DL1 with outcome in cancer and transplantation (Bari et al., 2013) (Babor et al., 2014) (Dutta, Saikia, Phookan, Baruah, & Baruah, 2014). However, some of the strongest associations come from disorders of pregnancy. For example, mothers with two *KIR A* haplotypes (*KIR AA* genotype) are at increased risk of disorders of placentation if the fetus carries a C2 epitope inherited from the father (Hiby et al., 2004; Nakimuli et al., 2015). Conversely, mothers with a *KIR B* haplotype (containing activating KIR2DS1 that can also bind C2) are at low risk, but instead these mothers have an increased risk of delivering a large baby (Hiby et al., 2014). When the fetus is homozygous for alleles encoding a C1 epitope (C1^+^HLA-C) the mother’s KIR genotype has no effect, so C2 is the crucial fetal ligand. These results suggest that binding of the maternal inhibitory KIR2DL1 to trophoblast C2^+^HLA-C increases the risk of pregnancy disorders, whereas activating KIR2DS1 promotes fetal growth. Functional experiments in mice support the idea that receptor/ligand interactions leading to strong NK inhibition impedes fetal growth (Kieckbusch, 2014, Gaynor & Colucci 2017, Moffett, & Colucci, 2014) and there is good evidence for natural selection against strong inhibitory C2-specific human *KIR2DL1* variants (Nemat-Gorgani, 2018).

Currently 26 *KIR2DL1* alleles have been identified which can be grouped according to which haplotypes they tend to segregate onto (Hilton, Guethlein, et al., 2015a). Those typically found on the *KIR A* haplotype in European populations are *KIR2DL1*003, *002* or *\*001*, denoted hereon as *KIR2DL1A*. The dominant allele on the *KIR B* haplotype is *KIR2DL1*004*, designated as *KIR2DL1B* (**Figure 1A**). These alleles vary in frequency across populations (**Figure 1A**) and show differences in expression levels of both RNA and protein (Babor et al., 2014; Dunphy et al., 2015; McErlean et al., 2010). Moreover, functional studies have shown that KIR2DL1 allotypes bind C2^+^HLA-C allotypes with variable affinities, form an hierarchy of inhibition in response to their cognate ligand and differ in their ability to respond to missing self (Bari et al., 2009; Hilton, Norman, et al., 2015b; Yawata et al., 2008). However, most of these studies have been restricted to using in vitro systems, cell lines or donors homozygous for the alleles of interest. Thus, it is unclear how these data translate into functional differences on primary NK cells. In particular, it is not known whether KIR2DL1 allotypes are co-dominantly expressed by NK cells or if they confer different educating signals when presented with identical HLA-C environments.

**Figure 1.**
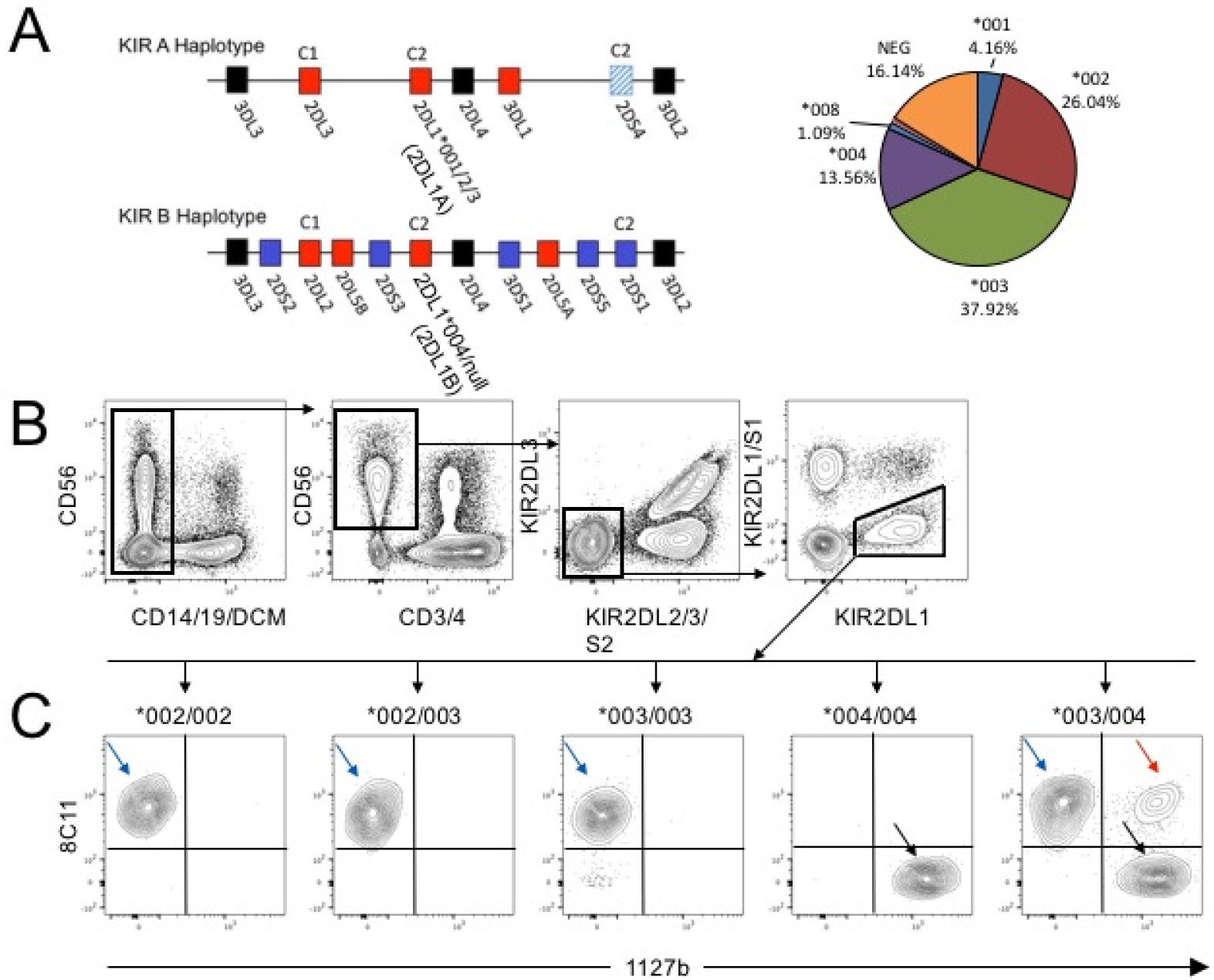
A new strategy to identify different KIR2DL1 allotypes in single NK cells. **A)** Representative KIR A and B Haplotypes. Cognate HLA-C ligands are depicted above their receptor. Framework genes are shown in black, activating KIR in blue and inhibitory KIR in red. In European populations the *KIR2DL1* alleles on the A haplotype are *KIR2DL1*001, *002* or *\*003*. The *KIR2DL1* allele on the B haplotype is almost exclusively *KIR2DL1*004*. Allele frequencies in the cohort in this study are depicted in the chart. **B)** Cryopreserved PBMCs from donors typed for *KIR2DL1* alleles were stained. Gating strategy to identify KIR2DL2/3/S2^neg^ 2DS1^neg^ 2DL1^pos^ NK cells in blood. **C)** Results for donors with different *KIR2DL1* alleles are shown. Three subsets were identified in donors heterozygous for *KIR2DL1*003* and *\*004***: \***003 single positive cells (003sp, blue arrow), *004 single positive cells (004sp, black arrows), and *003/*004 double positive cells (003004dp, red arrow).

We have identified a staining combination of anti-KIR antibodies that permits the separation of NK cells expressing KIR2DL1A and KIR2DL1B allotypes within the same individual. We find that KIR2DL1A allotypes are preferentially expressed by peripheral blood NK (pbNK) cells and KIR2DL1A^pos^ NK cells respond more strongly in missing self assays than KIR2DL1B^pos^ cells from the same individual and his or her HLA background. In a case-control study, we show that the risk of developing pre-eclampsia is associated with the number of *KIR2DL1A*, but not *KIR2DL1B* gene copy number. Finally, we study the expression of KIR2DL1 allotypes on the tissue resident population of NK cells that are likely to be driving this association in pregnancy. Similar to their peripheral blood counterparts, we find that in decidual NK cells, the KIR2DL1 positive niche is dominated by expression of KIR2DL1A allotypes.

## METHODS

### Cohorts and genotyping

Genomic DNA was obtained from three case/control UK Caucasian cohorts with pre-eclamptic patients and one prospective cohort: controls (n=696), pre-eclamptics (n=693). Informed written consent from the subjects and the corresponding ethical approvals: Cambridge Research Ethics Committee (reference number 01/197, 05/Q0108/367, 07/H0308/163), GOPEC (Derb reference number 09/H0401/66), London multiregional ethical committee (reference number) and Germany (reference number). Pre-eclampsia was defined by the clinical appearance of hypertension and proteinuria. For samples used for phenotyping and functional analysis, genomic DNA was isolated from decidual tissue and digested with proteinase K and RNase A (Roche) in combination with tissue lysis and protein precipitation buffers (Qiagen), prior to precipitation of DNA with isopropanol. For blood samples, Genomic DNA was isolated using the QIAamp DNA Mini Blood Kit (Qiagen). KIR genes presence/absence and HLA-C1/C2 genotyping for the three previously published pre-eclamptic cohorts and the families was performed by PCR-SSP as described previously (Hiby et al., 2004; 2010; 2008). The two remaining cohorts, the prospective cohort of controls and a cohort of pre-eclamptics, were typed for maternal KIR genes by multiplexed quantitative PCR (Jiang 2012). Genotyping of KIR2DL1 alleles was performed by pyrosequencing targeting exons 1, 4, 5, 7, and 9 (Norman 2013).

### KIR genotype analysis

KIR haplotype regions were defined following the KIR2011 Workshop recommendations. Briefly the centromeric A region (cA) was defined by the presence of KIR2DL3 and KIR2DL1, the centromeric B region (cB) by KIR2DS2 and KIR2DL2, the telomeric A region (tA) by KIR3DL1 and KIR2DS4, telomeric B region (tB) by KIR3DS1 and KIR2DS1.

### Primary tissue

Samples were obtained from three different sources. Firstly, previously characterized cryopresereved PBMCs from healthy donors were kindly provided by Karl-Johan Malmberg, Karolinska Institutet. Secondly, peripheral blood and matched decidua were obtained from women undergoing elective terminations of first trimester pregnancies. Mononuclear cells were isolated by enzymatic digestion of decidual tissue as described previously (Male, Gardner, & Moffett, 2001) and cryopreserved. Thirdly, peripheral blood was purchased from the NHS Blood and Transplant unit and PBMC were cryopreserved. Ethical approval for this study was obtained from the Cambridge Research Ethics Committee and Karolinska Institutet ethics review board with all participants supplying fully informed consent.

### Cell lines

YTS cells were stably transfected with a pcDNA3.1 plasmid encoding a single *KIR2DL1* allele (either *001, *003 or *004). These cells were obtained from Pippa Kennedy, Manchester University. Cells were grown in RPMI medium (10% FCS) under selection with G418 (1.5 mg/ml).

### Antibodies and flow cytometry

| Receptor | Clone | Fluorophore | Provider |
| --- | --- | --- | --- |
| CD3 | UCHT1 | PE-Cy5 | Beckman Coulter |
| CD4 | OKT4 | PE-Cy5 | BioLegend |
| CD14 | M5E2 | BV510 | Biosciences |
| CD19 | HIB19 | BV510 | Biosciences |
| CD56 | N901 | PE-Dazzle | BioLegend |
| KIR2DL1 | REA248 | PAC-Vio770 | Miltenyi |
| KIR2DL1/S1 | EB6 | PE-Cy7 | Beckman Coulter |
| KIR2DL2/3/S2 | GL183 | PE-Cy5.5 | Beckman Coulter |
|  | CH-L | BB515 | Biosciences |
| KIR2DL3 | 180701 | FITC | R&D Systems |
| KIR3DL1 | DX9 | BV421 | BioLegend |
| NKG2A | Z199 | APC | Beckman Coulter |
| KIR2DS1/L1 | 1127b | PE | R&D Systems |
| KIR2DL1/L2/3/S2 | 8C11 | Biotin | C. Retière |
| CD107a | H4A3 | Bv650 | BioLegend |

Antibodies were biotinylated using the Fluoreporter Mini-biotin-XX protein labelling kit (Life Technologies) and detected using streptavidin-Qdot 605 (Invitrogen/Thermo Fisher). Viability was assessed through staining with LIVE/DEAD Aqua (Invitrogen/Thermo Fisher). Cells were thawed in complete medium (RPMI 1640 medium, antibiotics, 10% Fetal Calf Serum), counted and distributed at approximately 1e6 cells/well in 200ul Facs Wash (FW, 1xPBS, 2% Fetal Calf Serum, 2mM EDTA). Cells were then stained for 1hr at 4C unless otherwise stated in 30ul of cocktail. KIR2DS1 was detected as previously described (Fauriat et al., 2010). After staining with a secondary cocktail containing LIVE/DEAD Aqua and streptavidin-Qdot 605 for 30 minutes at 4C, cells were fixed in 1% paraformaldehyde for 10 minutes at room temperature. Finally, fixed cells were resuspended in FW and analysed on a LSR Fortessa (BD Biosciences). The antibody 8C11 only bound KIR2DL1^pos^ cells in donors carrying the alleles *KIR2DL1*002* and/or *\*003*. The antibody 1127b only bound KIR2DL1^pos^ cells in donors carrying the allele *KIR2DL1*004* (**Figure 1**)

### Functional assays

PBMCs were thawed and rested overnight in complete medium. 5e5 PBMCs were co-cultured with 5e4 K562 target cells for 6 hours in 96-well U bottom plates at 37C. After 1 hour of incubation, Golgi Plug (BD Biosciences) and Golgi Stop (BD Biosciences) were added. This method is adapted from Bryceson et al 2010 (Bryceson et al., 2010). Cells were then stained and fixed as described above.

### Statistics

Categorical data was analysed using the chi-square and Fisher’s exact test with two-tailed mid-p adjustment and Student’s t-test for continuous data. The magnitude of the effect was estimated by odds ratios (OR) and their 95% confidence intervals (CI). For comparisons of matched groups, a repeated measures one-way ANOVA with Tukey’s multiple comparisons test was performed. Statistical analyses were largely performed using PRISM (GraphPad Software Inc.) and the open source statistical package R (www.r-project.org). A P-value of ≤ 0.05 was considered to be statistically significant.

## RESULTS

### A new strategy to identify different KIR2DL1 allotypes in single NK cells

One of the major stumbling blocks to studying the effects of allelic variation of KIR on NK cell function and its contribution to disease has been the lack of antibodies specific for individual KIR receptors. As a result, anti-KIR antibodies often display cross reactivity for the products of several KIR genes. This property has been exploited to study expression of different KIR2DL3 allotypes in heterozygotes, but has thus far not been possible for KIR2DL1 (Beziat et al., 2013; Falco et al., 2010). To address this, we transfected YTS cells with individual FLAG tagged *KIR2DL1A* (*KIR2DL1*001,003*) and *KIR2DL1B* (*KIR2DL1*004*) alleles. We then tested a panel of anti-KIR antibodies to determine their specificities (**Figure S1**). Two antibodies were identified that recognised selectively either KIR2DL1A allotypes, mAb 8C11, or KIR2DL1B, mAb 1127b. The binding profile for 8C11 was consistent with the predictions made by David et al. (David et al., 2009). Both antibodies also cross react with other KIR (**Table S1**) and thus we designed a panel that would be able to distinguish specific KIR2DL1 allotypes on primary NK cells. Cryopreserved PBMCs from donors typed for *KIR2DL1* alleles were then stained. *KIR2DL1*001* donors are lacking due to their rarity but, because, KIR2DL1*001 and *002 do not differ in their extracellular domains, results obtained for KIR2DL1*002 can be used as a proxy for KIR2DL1*001. Due to antibody cross reactivity, NK cells expressing KIR2DL2/3, KIR2DS2 and KIR2DS1 were gated out of the analysis (**Figure 1B**). As expected from our findings on YTS cells, 8C11 only recognises KIR2DL1A and not KIR2DL1B; conversely 1127b binds only KIR2DL1B and not KIR2DL1A allotypes (**Figure 1C**). As further confirmation, KIR2DL1A single positive (sp), KIR2DL1Bsp and KIR2DL1AB double positive (dp) subsets were sorted from a *KIR2DL1*003/004* heterozygous donor and *KIR2DL1* transcripts amplified by RT-PCR. The sequences of the *KIR2DL1* allele specific transcripts in each subset matched the protein data (data not shown). In addition, to ensure that there was no steric hindrance of the antibodies, we permuted the order of antibody staining (data not shown). For *KIR2DL1A/B* heterozygous donors, our gating strategy allowed us to measure the percentage of cells expressing each allotype in a subset of KIR2DL1 positive cells (KIR2DL2/3/S2^neg^,2DS1^neg^,2DL1^pos^ pbNK cells) (**Figure 2A**). Expression of KIR2DL1A dominates in all donors, with the mean proportion of KIR2DL1Asp pbNK cells observed being 56% in comparison to 35% for KIR2DL1B cells and only 8% for KIR2DL1ABdp cells (n = 20) (**Figure 2B,C**). These results show that, for the first time, we are able to visualise expression of two different *KIR2DL1* allotypes present on *KIR A* and *KIR B* haplotypes within the same heterozygous individual. Moreover, although there is co-dominant expression, KIR2DL1A is expressed by the majority of NK cells.

**Figure 2.**
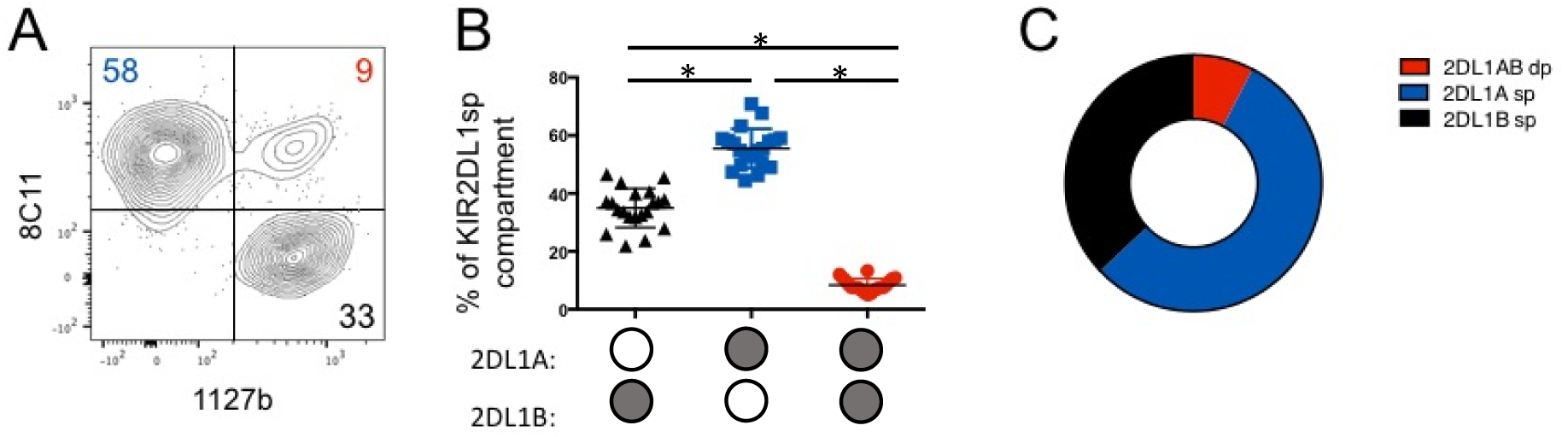
KIR2DL1A allotypes are preferentially expressed by KIR2DL1sp pbNK cells. Cryopreserved PBMCs from *KIR2DL1A/B* heterozygotes were stained as in Figure 1, (n=20). Cells were gated to identify KIR2DL2/3/S2^neg^ 2DS1^neg^ 2DL1^pos^ NK cells (KIR2DL1sp subset). **A)** KIR2DL1sp subset stained with 8C11 and 1127b in a donor heterozygous for *KIR2DL1A/B*. **B)** Proportions of KIR2DL1A single positive, KIR2DL1B single positive and KIR2DL1AB double positive subsets are shown as percentages of the KIR2DL1 subset summarised in the donut plot in panel **C**). * *P* <.0001; [RM one-way ANOVA, Tukey’s multiple comparisons test]

### KIR2DL1A single positive NK cells are more responsive than KIR2DL1B single positive NK cells

Although previous attempts to link *KIR2DL1* alleles to different NK functions have been limited, KIR2DL1*004 was found to be hypofunctional in the context of missing self (Yawata et al., 2008). However, this compared responses between donors with and without *KIR2DL1*004*, hence with different HLA backgrounds. We measured missing self responses of pbNK to the standard HLA class I-deficient K562 cell line to study how different KIR2DL1 allotypes within the same donor educate NK cells to alter their responsiveness. To rule out the influence of other inhibitory receptors, these were excluded using a flow cytometry gating strategy from individuals heterozygous for *KIR2DL1A* and *KIR2DL1B*. Thus, we focused on NKG2A^neg^ KIR2DL2/3/S2^neg^ KIR3DL1^neg^ KIR2DS1^neg^ KIR2DL1^pos^ NK cells (**Figure 3A**). Donors were further stratified by whether they possess a C2^+^HLA-C allele, the cognate ligand for KIR2DL1. Although both KIR2DL1A and KIR2DL1B are able to educate NK cells (**Figure 3B,C**), there is a significantly higher missing self response in KIR2DL1Asp compared with KIR2DL1Bsp NK cells. Moreover, an additive response in NK cells expressing both *KIR2DL1* alleles can be seen compared with KIR2DL1B but not KIR2DL1A. All three KIR2DL1 subsets were similarly hyporesponsive in C1/C1 donors. These results show that KIR2DL1A allotypes are better educators of NK cells than KIR2DL1B allotypes.

**Figure 3.**
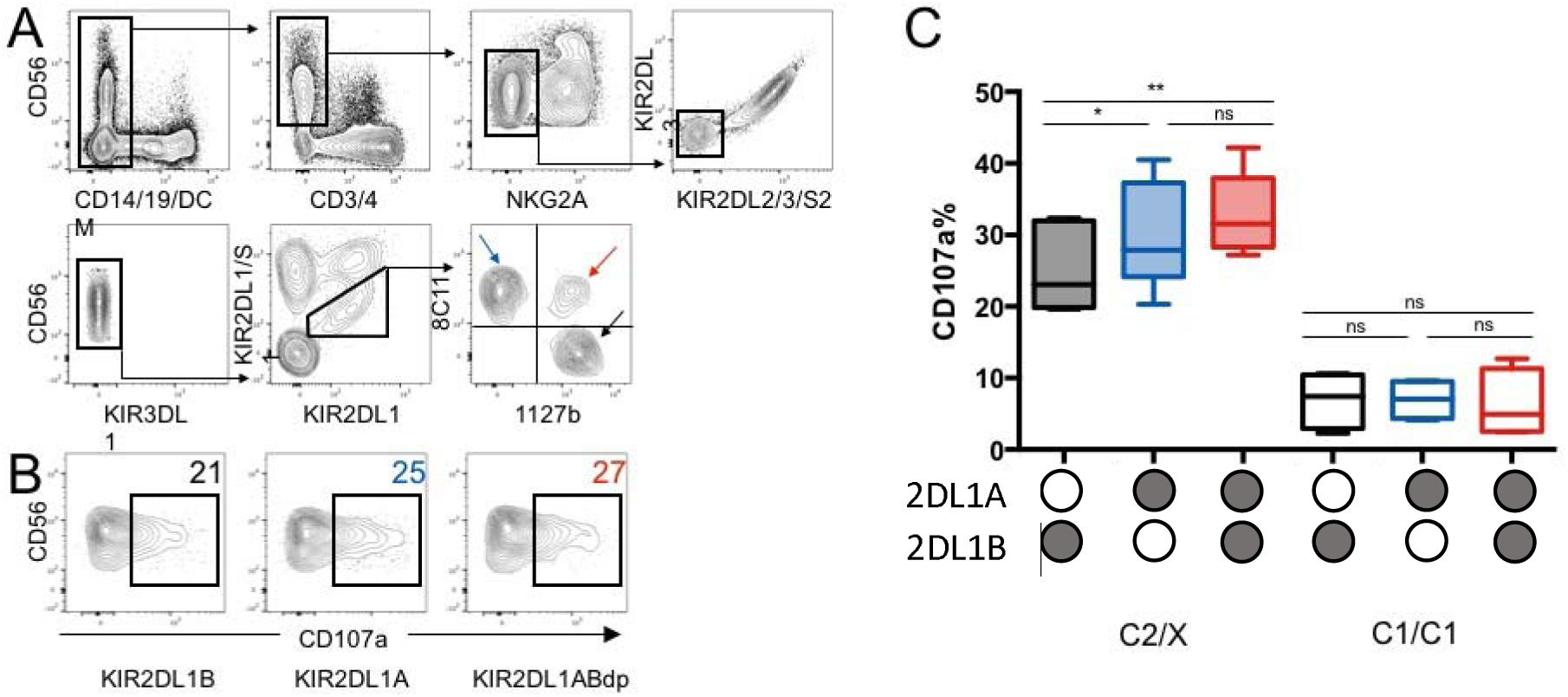
KIR2DL1A single positive NK cells are more responsive than KIR2DL1B single positive NK cells. Cryopreserved PBMCs were co-cultured with K562 and degranulation was assessed by staining with anti-CD107a antibody (n=12). All donors are heterozygous for *KIR2DL1A* and *B* alleles. **A)** NKG2A^neg^ KIR2DL2/3/S2^neg^ 3DL1^neg^ 2DS1^neg^ 2DL1^pos^ NK cells were identified. KIR2DL1Asp (blue arrow), KIR2DL1Bsp (black arrow) and KIR2DL1ABdp (red arrow)**; B)** Representative CD107a degranulation FACS plots for allele specific subsets from the same donor. From Left to Right: KIR2DL1Bsp, KIR2DL1Asp, KIR2DL1ABdp; **C)** Degranulation data for *KIR2DL1* allele specific subsets is shown for 12 donors. Donors are stratified by HLA-C status as mothers homozygous for *HLA-C* with C1 epitope (C1/C1) and those with at least one C2 epitope (C2/X). \**P* <.05; \*\**P* <.01; ns, not significant [RM one-way ANOVA, Tukey’s multiple comparisons test]

### KIR2DL1A allotypes and gene copy number associate with greater risk of pre-eclampsia

Having established that there are both phenotypic and functional differences between KIR2DL1 allotypes, we wanted to assess their impact in the context of risk of disease. Previous genetic studies have shown that mothers with two *KIR A* haplotypes are at increased risk of pre-eclampsia or fetal growth restriction, if the fetus carries a C2 epitope (Hiby et al., 2004). These results suggest that binding of KIR2DL1 to C2^+^HLA-C2 on placental extravillous trophoblast (ETV) cells leads to strong inhibition of decidual NK cells (dNK) and compromised placental development. To understand how different *KIR2DL1* alleles contribute to the risk of pregnancy disorders, we analysed a case/control cohort comparing 693 pre-eclamptic and 696 control pregnancies. When mothers lacking *KIR2DS1* are classified according to *KIR2DL1* genotype, the presence of *KIR2DL1A* was strongly associated with an increased risk of pre-eclampsia compared to mothers who lacked *KIR2DL1A* (P = 0.002, Odds Ratio (OR) = 2.2 [1.34-3.56]) (**Table 1**). This effect is seen even without correcting for fetal *HLA-C* genotype but is not evident in mothers who also had *KIR2DS1*, confirming an overriding protective effect of this activating KIR (**Table 1**).

**Table 1.** The presence of *KIR2DL1A* confers increased risk of PE

|  | Controls |  | Pre-eclamptic patients |  | P-value | OR | 95% CI |
| --- | --- | --- | --- | --- | --- | --- | --- |
|  | N | F(%) | N | F(%) |  |  |  |
| *KIR2DL1A* carriers | 598 | 85.9% | 628 | 90.6% | **0.007** | 1.58 | 1.14-2.21 |
| Others (no *KIR2DL1A*) | 98 | 14.1% | 65 | 9.4% |  |  |  |
| **KIR2DS1 negative** |  |  |  |  |  |  |  |
| *KIR2DL1A* carriers | 352 | 50.7% | 423 | 61.0% | **0.002** | 2.18 | 1.34-3.56 |
| Others (no *KIR2DL1A*) | 49 | 7.1% | 27 | 3.9% |  |  |  |
| **KIR2DS1 positive** |  |  |  |  |  |  |  |
| *KIR2DL1A* carriers | 245 | 35.3% | 205 | 29.6% | 0.751 | 1.08 | 0.68-1.71 |
| Others (no *KIR2DL1A*) | 49 | 7.1% | 38 | 5.5% |  |  |  |

Having identified that women who possess *KIR2DL1A* alleles are more at risk than those with other *KIR2DL1* genotypes we next addressed the effect of increasing the number of copies of *KIR2DL1A*. Mothers were grouped according to whether they had 0, 1 or 2 copies of *KIRDL1A* or *KIR2DL1B*. We found that increasing the copy number of *KIR2DL1A* was associated with increased risk of developing pre-eclampsia (P = 0.006). The increasing OR as *KIR2DL1A* copy number increases, indicates an additive genetic risk model (**Table 2**). The increased risk associated with *KIR2DL1A* copy number remained even when *KIR2DS1* was included as a covariate. In contrast, increasing copy number of *KIR2DL1B* had the opposite effect, lowering the risk of pre-eclampsia as *KIR2DL1B* copy number increased (OR, 0.68 [0.32-1.34]. Taken together, the results in Tables 1 and 2 show that *KIR2DL1A* alleles are associated with a greater risk of pre-eclampsia than *KIR2DL1B* alleles.

**Table 2.** KIR2DL1A confers more risk of PE as copy number increases

|  | Controls |  | Pre-eclamptic patients |  | P-value | OR | 95% CI |
| --- | --- | --- | --- | --- | --- | --- | --- |
|  | N | F(%) | N | F(%) |  |  |  |
| **KIR2DL1A CNV** |  |  |  |  |  |  |  |
| 0 | 135 | 10.6% | 56 | 7.8% | **0.006** | 1 |  |
| 1 | 539 | 42.3% | 282 | 39.4% |  | 1.26 | 0.90-1.79 |
| 2 | 599 | 47.1% | 377 | 52.7% |  | 1.52 | 1.09-2.14 |
| **KIR2DL1B CNV** |  |  |  |  |  |  |  |
| 0 | 961 | 75.5% | 553 | 77.3% | 0.261 | 1 |  |
| 1 | 284 | 22.3% | 151 | 21.1% |  | 0.92 | 0.74-1.15 |
| 2 | 28 | 2.2% | 11 | 1.5% |  | 0.68 | 0.32-1.34 |

CNV is copy number variation of KIR2DL1A or KIR2DL1B alleles. The value can be 0, 1 or 2 in this regression analysis.

### Preferential expression of KIR2DL1A allotypes in tissue resident decidual NK cells

Our findings show that *KIR2DL1A* compared to *KIR2DL1B* alleles are associated with the risk of developing pre-eclampsia and with both phenotypic and functional differences in pbNK cells. However, it is decidual NK (dNK) cells, present in the lining of the pregnant uterus (decidua), rather than pbNK that are likely to be responsible for the biological effects on placental development. Early in pregnancy, dNK cells account for ~70% of the decidual leukocytes (King, Balendran, Wooding, Carter, & Loke, 1991). They interact with surrounding maternal cells in the decidua and with the invading fetal trophoblast cells that express HLA-C molecules. The receptor expression profiles of dNK cells are quite different than pbNK cells (Ivarsson et al., 2017; Male et al., 2011) with a greater proportion of dNK cells expressing KIR2DL1 compared to pbNK cells from the same donor (Xiong et al., 2013). Education of decidual NK cells is also regulated differently from that of pbNK cells (Sharkey et al., 2015). We compared the expression of KIR2DL1A and KIR2DL1B allotypes on dNK cells in heterozygous individuals in a similar analysis to Figures 1 and 2. dNK cells express both KIR2DL1A and KIR2DL1B allotypes and co-dominant expression of both allotypes is observed in heterozygotes (**Figure 4A, 4B**). In accordance with the results of pbNK cells (**Figure 2**), for *KIR2DL1A/B* heterozygous donors, there is preferential expression of KIR2DL1A allotypes (57%) in KIR2DL1sp dNK cells (n= 7) (**Figure 4C**) compared to 34% for KIR2DL1Bsp and 7% for KIR2DL1ABdp. When comparing matched NK cells from blood and decidua, the expression patterns of the KIR2DL1 allotypes were very similar (**Figure S2**). Taken together these results indicate that the mechanism governing the relative frequency of *KIR2DL1A/B* allele expression is similar in pbNK cells and in dNK cells.

**Figure 4.**
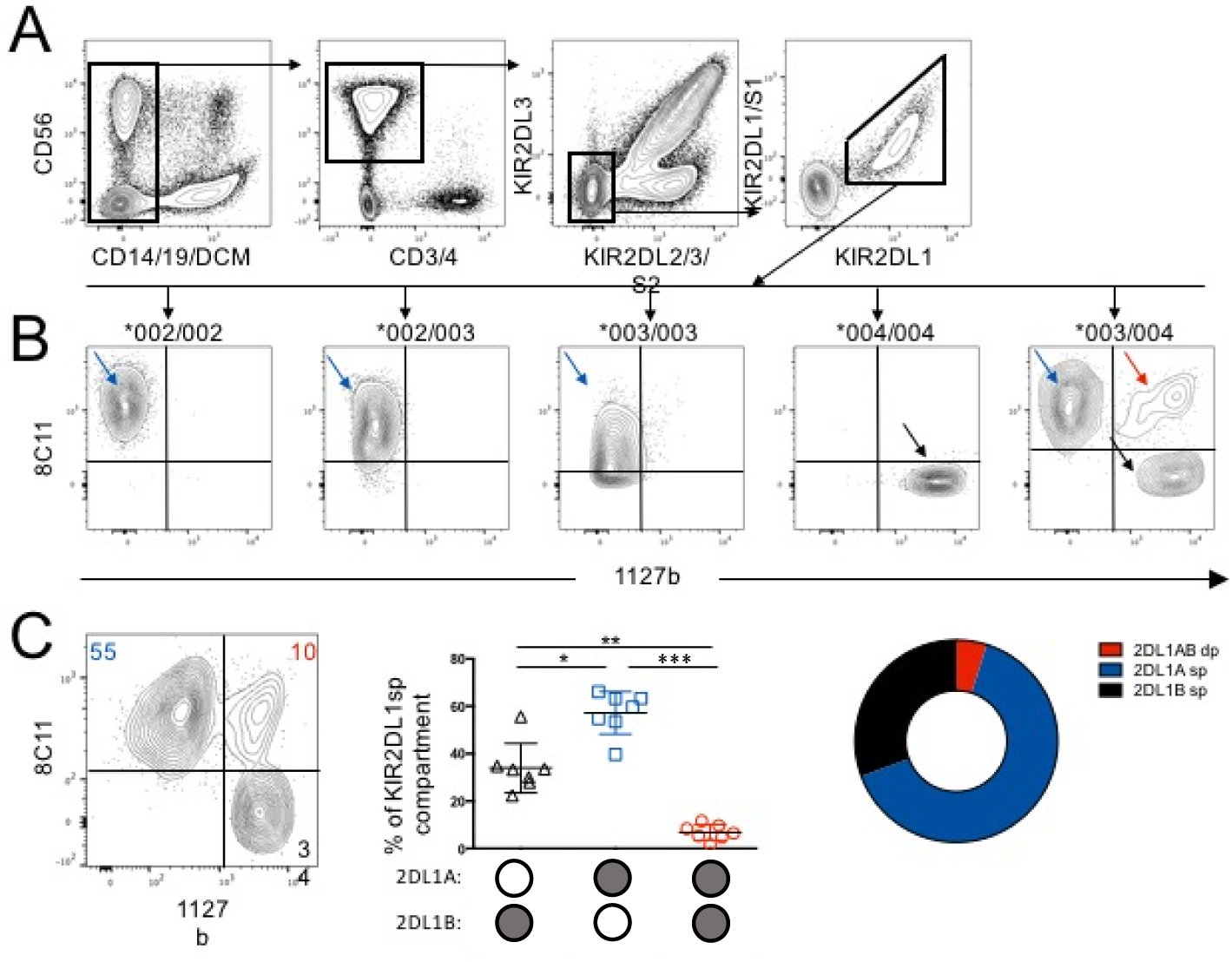
Preferential expression of KIR2DL1A allotypes in tissue resident decidual NK cells. Cryopreserved decidual mononuclear cells from donors typed for *KIR2DL1* alleles were stained to identify dNK cells expressing KIR2DL1 allotypes (n=7). **A)** Gating strategy to identify KIR2DL2/3/S2^neg^ 2DS1^neg^ 2DL1^pos^ dNK cells; **B)** dNK cells from donors typed for *KIR2DL1* alleles were stained and gated as in (A). Results for donors with different *KIR2DL1* alleles are shown above each chart; **C)** 2D FACS plot is representative of biallelic KIR2DL1 expression in dNK cells from a donor heterozygous for *KIR2DL1*003* and *\*004*. Proportions of KIR2DL1A single positive, KIR2DL1B single positive and KIR2DL1AB double positive subsets are shown as percentages of the KIR2DL1 subset and summarised in the donut plot. \**P* <.05; \*\**P* <.01; *** *P* <.0001; ns, not significant [RM one-way ANOVA, Tukey’s multiple comparisons test]

## DISCUSSION

We have developed a method to identify allotype-specific NK-cell subsets for the polymorphic receptor KIR2DL1. Using this method, we show that alleles associated with either *KIR* haplotype, *KIR2DL1A* and *KIR2DL1B*, are both expressed in heterozygous individuals, with *KIR2DL1A* prevailing both phenotypically and functionally in both peripheral and uterine NK cells. Because *KIR2DL1A* are associated with an increased risk to develop pre-eclampsia, this result could help to focus on patients at higher risk.

The lack of KIR specific antibodies has limited studies of individual KIR. This problem is amplified when studying *KIR* alleles because alleles can differ by a handful of residues. More recently, by combining cross-reactive anti-KIR antibodies it has been possible to achieve improved resolution. For example, KIR2DL3*005 allotypes can be identified from other KIR2DL3 allotypes using an unexpected cross reactivity of the anti-KIR2DL1/S1 clone, EB6 (Falco et al., 2010). By exploiting the cross reactivity of the anti-KIR antibodies, 8C11 and 1127b (David et al., 2009), we are able to visualise expression of two different *KIR2DL1* alleles carried on *KIR A* and *KIR B* haplotypes within the same heterozygous individual.

The surface expression of a particular KIR can be affected by allelic polymorphism, promoter differences, copy number variation, HLA ligand background and HCMV status (Béziat et al., 2012; 2013; Dunphy et al., 2015; Gardiner et al., 2001; Li, Pascal, Martin, Carrington, & Anderson, 2008). In donors homozygous for *KIR2DL1* alleles, the most common *KIR2DL1B* allele in Europeans, *KIR2DL1*004*, is expressed by significantly fewer NK cells than the most common *KIR2DL1A* allele, *KIR2DL1*003* (Dunphy et al., 2015). In accordance with this, and although there is bi-allelic expression of KIR2DL1 allotypes, KIR2DL1Asp NK cells dominated the KIR2DL1sp NK cell niche in our *KIR2DL1A/B* heterozygous donors.

Using this approach, the effects of KIR2DL1 alleles on NK function can, for the first time, be compared within the same individual. In this scenario, NK cells expressing different KIR2DL1 allotypes have been educated in identical HLA class I environments. By reducing variation attributed to inter-individual differences we have been able to assess the functional capabilities of KIR2DL1 allotypes. KIR2DL1A proved to be the strongest educator when compared to KIR2DL1B in missing self assays, in agreement with previous data which shows *KIR2DL1*004* to be hyporesponsive in comparison to KIR2DL1 alleles in other donors (Yawata et al., 2008). In contrast to education by KIR2DL3 alleles which was found to be non-additive with respect to degranulation (Beziat et al., 2013), our data suggests that there is an additive effect, with a significant difference in responsiveness between cells expressing KIR2DL1B alone, compared to KIR2DL1ABdp pbNK cells in donors with the C2 epitope that binds both KIR2DL1A and KIR2DL1B allotypes.

From our data it is not clear whether the differences in function of these alleles are attributable to differences in ligand binding avidity or in the quality or quantity of signalling. Binding of Fc fusion proteins of KIR2DL1A allotypes to microbeads coated with C2^+^HLA-C is only slightly higher than for KIR2DL1*004 Fc (Hilton, Norman, et al., 2015b). Notably, the significant differences in binding of each KIR2DL1 Fc protein to different C2^+^HLAC allotypes, emphasises the importance of comparing functional effects of *KIR2DL1* alleles in the same HLA-C background. Following ligand binding, KIR2DL1 proteins also differ in their signalling capacity with an arginine residue at position 245 (only in KIR2DL1A European alleles) in the transmembrane domain integral to the quality of signalling and durability of surface expression (Bari et al., 2009).

Genes encoding KIR receptors and their HLA ligands are associated with pregnancy disorders. Homozygosity for the *KIR* A haplotype (KIR AA) in the mother is more frequently associated with the disorders known as the Great Obstetrical Syndromes (pre-eclampsia, recurrent miscarriage and fetal growth restriction) (Hiby et al., 2004; 2008; 2010; 2014). We now show that *KIR2DL1A* but not *KIR2DL1B* increases the risk of developing pre-eclampsia in a dose dependent manner, particularly when *KIR2DS1*, the antagonist for *KIR2DL1*, is absent. The only precendent for allele specific associations with pregnancy disorders is confined to an activating KIR. Certain African *KIR2DS5* alleles are associated with decreased risk of pre-eclampsia (Nakimuli et al., 2015). Our finding of an association of specific KIR2DL1 alleles in pre-eclampsia is the first report of allele specific effects for an inhibitory KIR on pregnancy outcome.

To further understand how allelic variants can drive disease progression we analysed the NK cell population that is physiologically relevant to pre-eclampsia, namely dNK cells and show that like the blood compartment, KIR2DL1A is preferentially expressed by KIR2DL1sp dNK cells. Alleles of a gene coding for an inhibitory KIR may mediate their effects via two different mechanisms – NK-cell education and inhibition. Through calibrating the functional potential of dNK cells, education by the mother’s HLA-C could affect the ability of dNK cells to regulate trophoblast invasion. For example, genetic evidence suggests that for pregnancies in which the fetus is *C2*^+^*HLA-C*, mothers who are *KIRAA* and *C2*^+^*HLA-*C are at lower risk of developing pre-eclampsia than *KIRAA and C1C1*^+^*HLA-C* mothers (Hiby et al., 2010). This is consistent with the idea that maternal KIR2DL1^pos^ dNK cells are educated by C2^+^HLA-C prior to pregnancy. Alternatively, if excessive NK cell inhibition occurs through *KIRAA* dNK cells and fetal *C2 HLA-*C on invading trophoblast during pregnancy, this may lead to reduced remodelling of the uterine vasculature and poorer pregnancy outcomes in both humans and mice (reviewed in (Moffett & Colucci, 2014)) (Kieckbusch et al., 2014). There exists a hierarchy of inhibition which places common KIR2DL1A above KIR2DL1B in response to 721.221 cells expressing the *C2 HLA-C* allele *Cw6* (Bari et al., 2009). This model would predict increased inhibition when dNK expressing KIR2DL1A encounters fetal trophoblast expressing C2^+^HLA-C compared to dNK expressing KIR2DL1B. The ability to stain both KIR2DL1 allotypes within NK cells from the same genetic background will, for the first time, permit dissection of both the effects of maternal HLA-C on dNK education, and whether KIR2DL1 allotypes generate different responses to fetal HLA-C.

Genetic studies that analyse KIR allelic variation are increasingly common (Ahn et al., 2016; Bari et al., 2013; Martin et al., 2007; Nakimuli et al., 2015; Nemat-Gorgani et al., 2018). Of equal importance, is the need to develop methods that allow us to characterise how these allelic variants contribute to functional differences at the cellular level and ultimately cause pathology. Allelic variability of other KIR genes such as *KIR3DL1, KIR2DL2/3* and *KIR3DL2* has an impact on phenotype and function in the context of infection and autoimmunity (Augusto et al., 2015; Bari et al., 2016; Khakoo et al., 2004). For example, *KIR3DL1* polymorphisms associate with delayed HIV progression and predict patient survival in response to immunotherapy for neuroblastoma(Forlenza et al., 2016; Martin et al., 2007). We have developed a method that will allow assessment of phenotype and function of KIR2DL1 allele-specific subpopulations of cells within an individual. This will facilitate our understanding of how KIR2DL1 receptor polymorphisms affect cellular function and ultimately contribute to disease progression.

## Acknowledgements

We thank the members of the Colucci and the Moffett labs for suggestions and discussions. This work was funded by MedImmune, the Wellcome Trust (Grant 200841/Z/16/Z to FC and AM), the Medical Research Council to AS, the Centre For Trophoblast Research (CTR) and the Cambridge NIHR BRC Cell Phenotyping Hub.

We thank Karl-Johan Malmberg for providing PBMCs from healthy donors.

OH was supported by a MedImmune-Cambridge PhD fellowship.

OC was supported by a CTR Next Generation Fellowship

## Authors’ contribution

OH designed and executed experiments, analysed data and wrote manuscript

OC designed and executed experiments, analysed data and wrote manuscript

MI designed experiments, analysed data and edited manuscript

CR provided resources

TV executed experiments

HG provided resources and edited manuscript

AM designed experiments, analysed data and edited manuscript

AS designed experiments, analysed data and wrote manuscript

FC designed experiments, analysed data and wrote manuscript

## SUPPLEMENTARY FIGURES and TABLE

**Figure S1.**
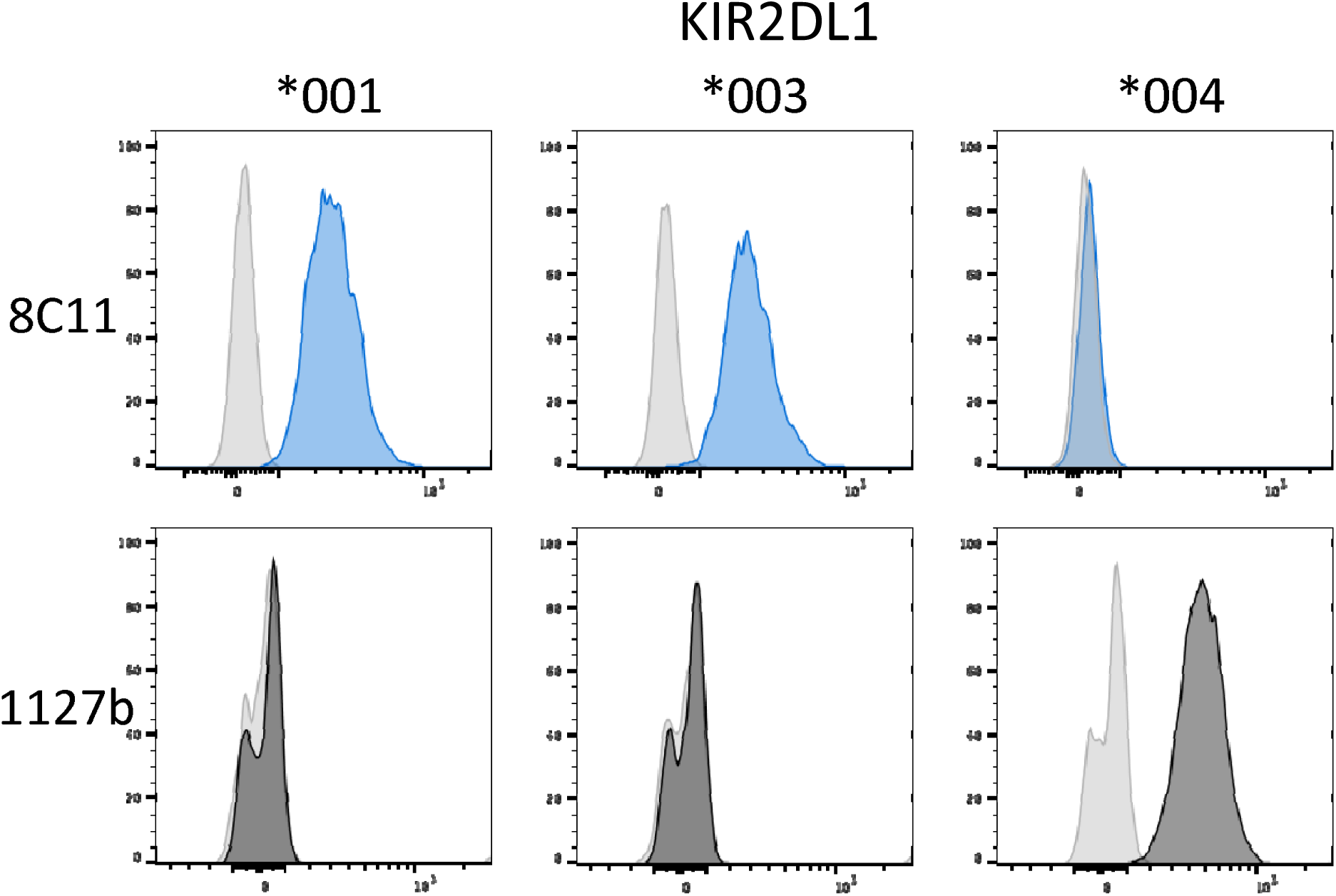
Antibody binding profiles of YTS cells transfected with *KIR2DL1* allotypes. YTS cells transfected with *KIR2DL1*001, *003* and *\*004* were stained with 8C11 and 1127b. The IgG control is shown in grey, 8C11 staining in blue and 1127b staining in black.

**Figure S2.**
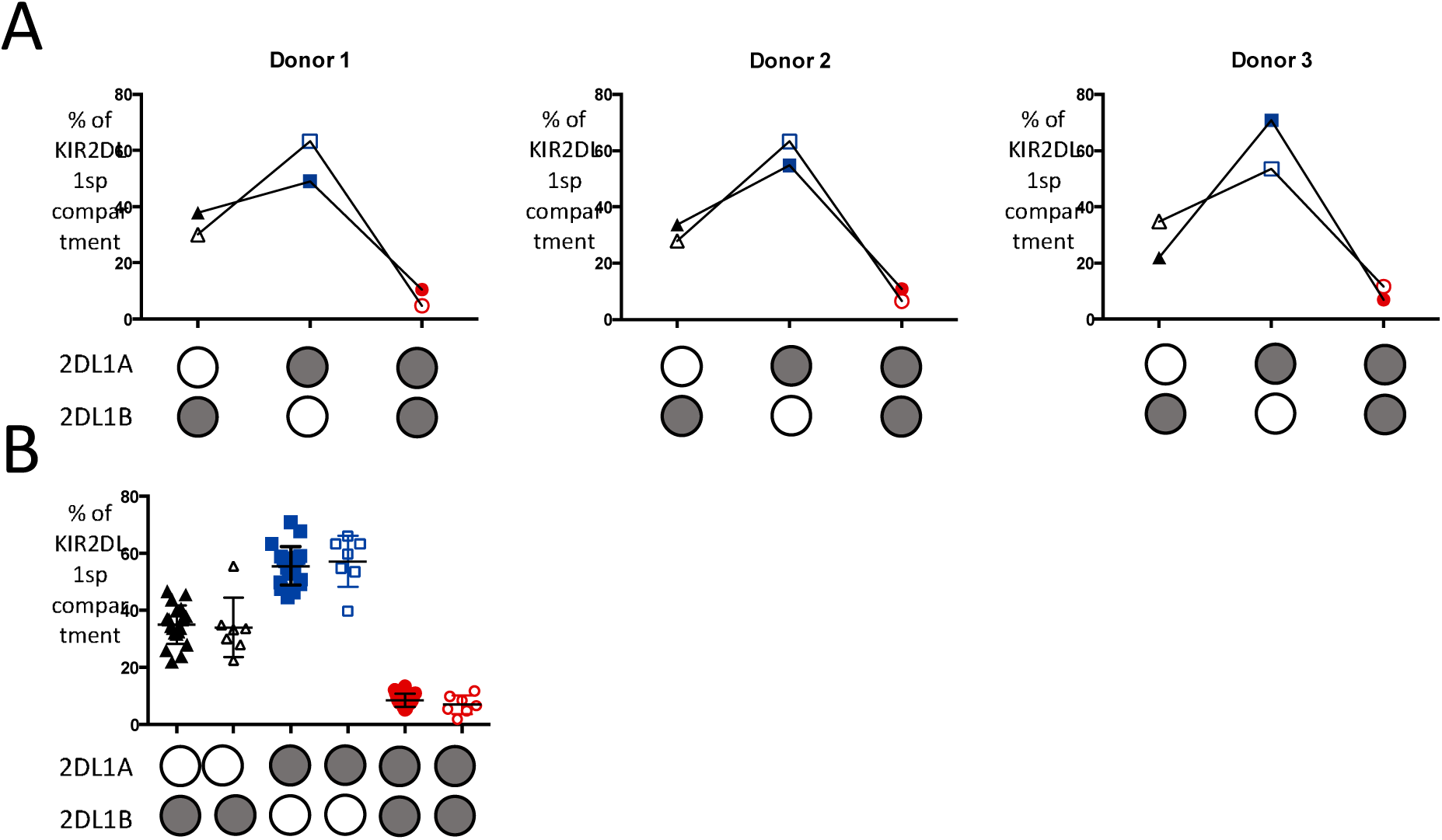
KIR2DL1 allotype expression is similar between peripheral blood and decidua. Cryopreserved decidual and peripheral blood mononuclear cells from donors heterozygous for *KIR2DL1A* and *KIR2DL1B* were stained. KIR2DL2/3/S2^neg^ 2DS1^neg^ 2DL1^pos^ NK cells were identified and proportions of KIR2DL1A^pos^ and KIR2DL1B^pos^ cells calculated. Data from 3 matched pairs **(A)** and all *KIR2DL1A/B* heterozygotes **(B)** are shown. Filled shapes = peripheral blood, empty = decidual.

**Table S1.**
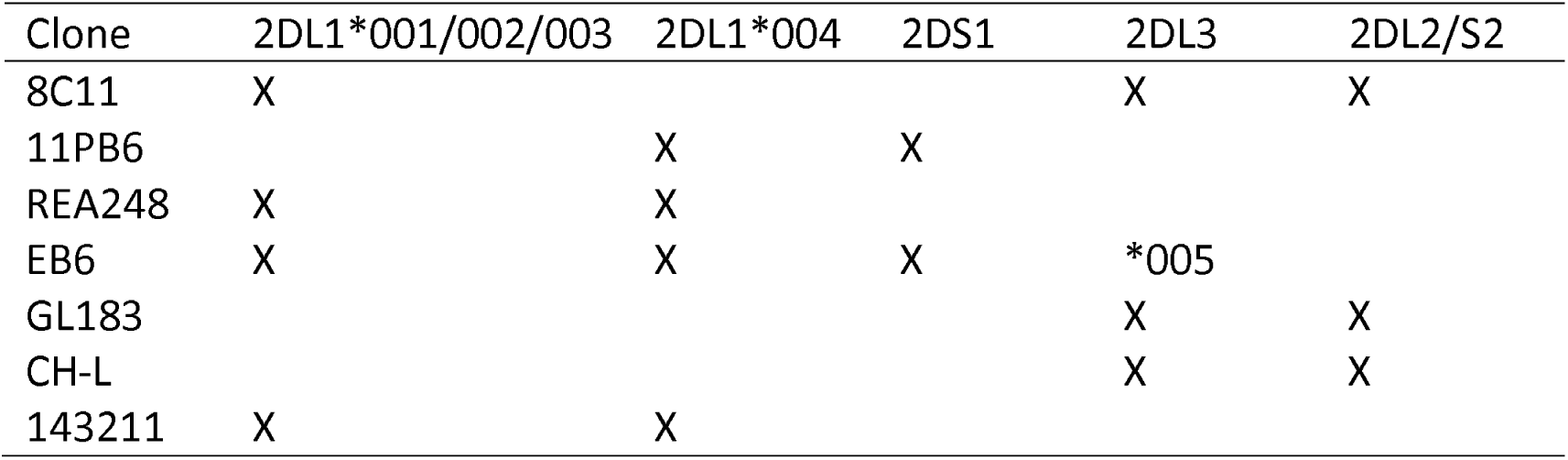
Cross-reactivity of KIR-specific monoclonal antibodies, which are indicated by their clone name

